# Ultraconserved elements help resolve the phylogeny of an ancient radiation of venomous flies (Diptera: Asilidae)

**DOI:** 10.1101/2020.11.09.375196

**Authors:** Chris M. Cohen, Katherine Noble, T. Jeffrey Cole, Michael S. Brewer

## Abstract

Robber flies or assassin flies (Diptera: Asilidae) are a diverse family of venomous predators. The most recent classification organizes Asilidae into 14 subfamilies based on a comprehensive morphological phylogeny, but many of these have not been supported in a subsequent molecular study using traditional molecular markers. To address questions of monophyly in Asilidae, we leveraged the recently developed Diptera-wide UCE baitset to compile seven datasets comprising 151 robber flies and 146 - 2,508 loci, varying in the extent of missing data. We also studied the behavior of different nodal support metrics, as the non-parametric bootstrap is known to perform poorly with large genomic datasets. Our ML phylogeny was fully resolved and well-supported, but partially incongruent with the coalescent phylogeny. Further examination of the datasets suggested the possibility that GC bias had influenced gene tree inference and subsequent species tree analysis. The subfamilies Brachyrhopalinae, Dasypogoninae, Dioctriinae, Stenopogoninae, Tillobromatinae, Trigonomiminae, and Willistonininae were not recovered as monophyletic in either analysis, consistent with a previous molecular study. The inter-subfamily relationships are summarized as follows: Laphriinae and Dioctriinae (in part) are successively sister to the remaining subfamilies, which form two clades; the first consists of a grade of Stenopogoninae (in part), Willistonininae (in part), Bathypogoninae+Phellinae, Stichopogoninae, Leptogastrinae, Ommatiinae, and Asilinae; the second clade consists of a thoroughly paraphyletic assemblage of genera from Dioctriinae (in part), Trigonomiminae, Stenopogoninae (in part), Tillobromatinae, Brachyrhopalinae, and Dasypogoninae. We find that nodal support does not significantly vary with missing data. Furthermore, the bootstrap appears to overestimate nodal support, as has been reported from many recent studies. Gene concordance and site concordance factors seem to perform better, but may actually underestimate support. We instead recommend quartet concordance as a more appropriate estimator of nodal support. Our comprehensive phylogeny demonstrates that the higher classification of Asilidae is far from settled, and it will provide a much-needed foundation for a thorough revision of the subfamily classification.

## Introduction

Robber flies or assassin flies (Diptera: Asilidae) are a diverse family of venomous predators (e.g., Figure 1) that likely originated in the Lower Cretaceous, ∼128 mya (Dikow *et al*., 2017). The most recent classification organizes Asilidae into 14 subfamilies and is based on the most comprehensive family-level phylogeny to date with 158 assassin fly species and 220 morphological characters analyzed in a parsimony framework (Dikow, 2009a; but see Karl, 1959 & Bybee *et al*., 2003). However, at least six of the new or revised subfamily concepts cannot be readily identified using external morphology, and consequently, 100 genera not examined by Dikow are currently without subfamily assignments or are otherwise *incertae sedis*. In addition, molecular evidence to date does not fully support this classification. A subsequent total-evidence analysis, which included 77 assassin flies, five genes, and 211 morphological characters, only recovered half of the twelve included subfamilies as monophyletic (Dikow, 2009b). The subfamilies Stenopogoninae, Dasypogoninae, Brachyrhopalinae, Willistonininae, and Tillobromatinae *sensu* Dikow are particularly problematic, not being supported with molecular data.

**Fig 1.**
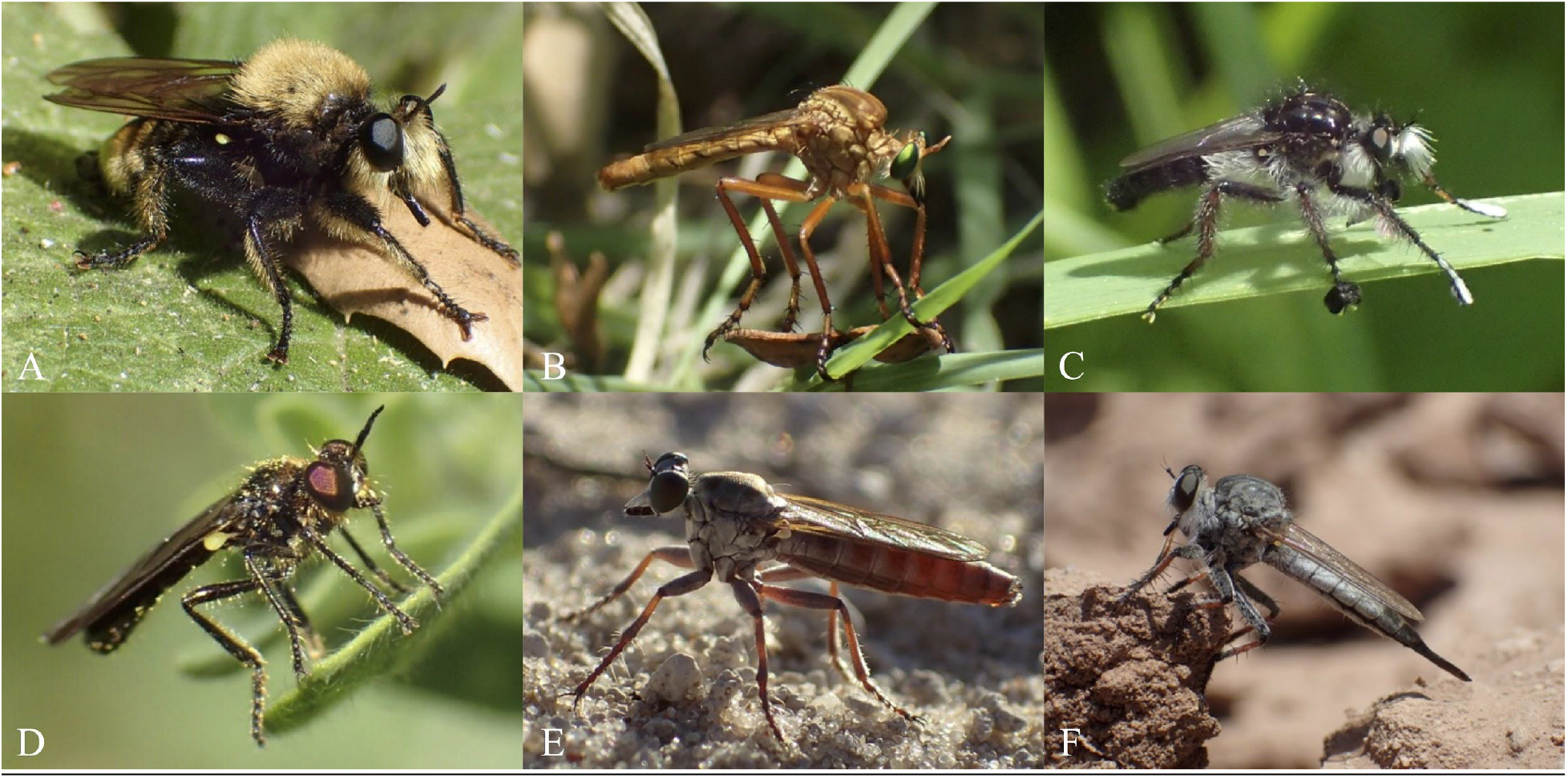
Photographs of select representatives of Asilidae. (A) *Laphria sp*. (Laphriinae). (B) *Diogmites sp*. (Dasypogoninae). (C) *Cyrtopogon sp*. (Brachyrhopalinae). (D) *Metadioctria resplendens* (Dioctriinae). (E) *Stichopogon sp*. (Stichopogoninae). (F) *Efferia sp*. (Asilinae). All photos by C.M. Cohen.

Both morphology and Sanger sequencing-based methods utilizing relatively few genes have thus been inadequate for resolving deep relationships within Asilidae. Phylogenomic approaches applied to assassin flies may alleviate these issues (and in Diptera generally; see Shin *et al*., 2018). Dikow *et al*. (2017) published Maximum Likelihood and ASTRAL phylogenies inferred using over 9,000 transcriptome-derived orthologs and nine assassin fly species. However, this shallow taxon sampling is insufficient for inferring accurate higher-level relationships. Phylogenomic approaches that allow for the cost-effective sequencing of many taxa have recently been developed. For example, Ultraconserved elements (UCEs) provide orders of magnitude more data at lower costs per-specimen - and data with better phylogenetic signal - than traditional sanger-based datasets (e.g., Blaimer *et al*., 2015; Gilbert *et al*., 2015; Zhang *et al*., 2019), are much cheaper per-specimen than transcriptome-based methods, and are capable of recovering phylogenetic loci from museum specimens (Blaimer *et al*., 2016; Van Dam *et al*., 2017). UCEs are useful for recovering both deep (Faircloth *et al*., 2012; Starrett *et al*., 2016; Branstetter *et al*., 2017; Baca *et al*., 2017; Buenaventura *et al*. 2020) and shallow relationships (Smith *et al*., 2014; Harvey *et al*., 2016; Manthey *et al*., 2016). The recently developed Diptera-wide UCE baitset (2.7kv1; Faircloth, 2017) was successfully tested in a group of closely related genera in the Empididae (Rhodén & Wahlberg 2020). We evaluate the utility of this baitset in reconstructing family-level relationships by combining it with comprehensive taxon sampling of the family Asilidae. The well-resolved and well-supported phylogeny of robber flies provided here will allow for the long-needed revision of the subfamily classification as well as provide an evolutionary framework for studies in comparative biology.

## Materials & Methods

### Taxon Sampling

Well-informed and carefully considered taxon sampling can ameliorate long-branch attraction (Heath, Hedtke, & Hillis, 2008), help resolve difficult nodes (Hillis, 1998), and increase overall phylogenetic accuracy (Zwickl & Hillis, 2002). Regarding ingroup sampling, 151 robber flies specimens representing 139 genera from all 14 subfamilies (*sensu* Dikow, 2009a) were included in phylogenetic analyses. This sampling encompasses all zoogeographic regions with particular emphasis on Nearctic and Australian taxa. Nine outgroup taxa were included, representing the closely related asiloid families Apioceridae and Mydidae, as well as a more distant outgroup, Bombyliidae (see Table S1 for a list of all included taxa). Freshly-collected specimens were preserved in 95% EtOH and stored in a -20°C freezer until ready for DNA extraction. Ethanol- or liquid nitrogen-preserved specimens were also borrowed from the Queensland Museum, Insect Genomics Collection at Brigham Young University, the Global Genome Initiative at the Smithsonian Institution, and the private collection of Dr. Darren Pollock.

Specimens were examined and identified by the first author. In cases where the genus or species was not immediately known, such specimens were identified to genus using available regional keys (e.g., Wood, 1981; Fisher, 2009; Londt & Dikow, 2017; Papavero, Artigas & Lamas, 2009) and to species using relevant keys from the literature (e.g., Barnes, 2008; Daniels, 1976; Martin, 1959; Paramonov, 1958). Specimens were photographed in ethanol using a Canon 6D camera and Canon 100mm and MP-E 65mm lens integrated with the BK Lab Imaging System (Dun, Inc.) prior to tissue extraction, whenever possible. This preserves a morphological record of the specimen and will aid in reconfirmation of identifications if necessary.

### Probes

The Diptera 2.7Kv1 baitset developed by Faircloth (2017), which uses 31,328 baits to target 2,711 loci, was chosen for this study in order to test its utility for resolving family-level relationships in Diptera. This probe set was generated using genomes from *Tipula oleracea, Aedes aegypti, Lutzomyia longipalpis*, and *Mayetiola destructor* (all “lower Diptera” = “Nematocera”), as well as *Megaselia abdita, Drosophila melanogaster*, and *Musca domestica* (all “higher Diptera” = Cyclorrhapha) (Faircloth, 2017, supplemental). No genomes from Asilidae or close relatives (i.e., lower Brachycera or “Orthorrhapha”) were included in probe development (see Wiegmann *et al*., 2011 and Shin *et al*., 2018 for discussions of higher taxa in Diptera).

### DNA Extraction and Sonication

Genomic DNA was extracted from leg and/or thoracic muscle tissue using the DNeasy Blood and Tissue Kit (Qiagen, Valencia, CA) following the manufacturer’s protocol. A Qubit 2.0 fluorometer (Life Technologies) was used to determine DNA concentration. Select extractions were run on a gel to determine shearing time and evaluate shearing success. Most material was recently collected and well-preserved, so standard shearing parameters were used for all samples. Samples were sheared using a Qsonica sonicator (Q800R1; Qsonica, LLC) to achieve an average fragment size of 600 bp.

### Library Preparation

Libraries were prepared with the KAPA Hyper Prep Kit (Kapa Biosystems) following Faircloth *et al*. (2014), but using a modified protocol developed by Michael Branstetter for use with the iTru dual-indexing adapter system (Glenn *et al*., 2016). The iTru system uses two 8 bp indexes for each sample, allowing for greater multiplexing and reducing adapter contamination. Briefly, this protocol consists of steps to first bind DNA to magnetic beads, then washed with EtOH. DNA is then subjected to a blunt end repair + A-tailing reaction that prevents chimeric formation. This is followed by ligation of the Y-yoke adapters and another bead binding and EtOH wash. A PCR reaction with i5 and i7 index primers is then performed, followed by additional bead washes. The final DNA elution was quantified in a Qubit to verify library success and to prepare for pooling.

### Pooling, Enrichment & Sequencing

First, 1-10 libraries were pooled together at equimolar ratios, producing 10 pooled libraries for 96 samples. These pooled libraries were then enriched using the myBaits custom kit for Diptera (Arbor Biosciences, Inc.) following the protocol developed by Faircloth *et al*. (2014) and modified by Michael Branstetter and Katherine Noble. Briefly, a blocking mix is added to the adapter-ligated DNA, preventing them from hybridizing. The probes are then added and non-target DNA is washed away, thus producing libraries enriched for Diptera UCEs. The DNA concentration for the enriched libraries were determined using Qubit, and qPCR was used to verify enrichment success. Enriched libraries were multiplexed and sequenced across multiple lanes of Illumina HiSeq 2500 at the High-Throughput Genomics Center at the University of Utah and and Illumina NovaSeq 6000 at HudsonAlpha. Raw reads were deposited in the NCBI Sequence Read Archive (SRA; Bioproject accession number PRJNA666700).

### Processing

Raw reads were processed with PHYLUCE v1.6.5 (Faircloth, 2017) and Illumiprocessor v2.0.9 (Faircloth, 2013). Resulting processed reads were assembled in Trinity v2.1.1 (Grabherr *et al*., 2011). PHYLUCE was used to match the resulting contigs to the probes using the script phyluce_assembly_match_contigs_to_probes. An additional five taxa were represented by transcriptomes and one from a genome. UCE loci were extracted from these assemblies using the scripts phyluce_probe_run_multiple_lastzs_sqlite (identity and coverage set to 0.65) and phyluce_probe_slice_sequence_from_genomes in PHYLUCE. The resulting FASTA files were added to the “contig” directory containing the other UCE assemblies. After extracting the UCE loci with the scripts phyluce_assembly_get_match_counts and phyluce_assembly_get_fastas_from_match_counts, individual orthologs were aligned with mafft v.7.313 (Katoh & Standley, 2013) and trimmed with trimAL v1.4.rev15 (Capella-Gutierrez, Silla-Martinez, & Gabaldon, 2009).

Screening for site homology is an important consideration, as problems with sequence alignment can have negative effects on topology and branch lengths, even for phylogenomic datasets (Springer & Gatesy, 2018). However, due to the scale of the dataset, individual alignments were not manually inspected. Instead, the program Spruceup (v2019.2.3) was used to detect outlier sequences (Borowiec, 2019). Rather than simply removing problematic columns from the alignment like traditional trimming programs, Spruceup can detect and remove poorly-aligned rows or sequence fragments from alignments. All assembled and trimmed loci were concatenated into a single matrix and provided as input to Spruceup along with partition and config files. Default settings and no guide tree were used. After “un-concatenating” the trimmed output matrix entitled “0.95 lognorms cutoff” (the most conservative cutoff) with a custom script, some loci had so much sequence data trimmed from certain taxa that these loci needed to be removed from the alignments. Therefore a custom script was used to replace taxon sequences with insufficient characters (less than 10% length of alignment) with missing characters (“?”) for each alignment. Sequences composed entirely of missing characters were then manually removed from alignments, resulting in 11 alignments being dropped for having less than three taxa. The remaining alignments were again trimmed with trimAL v1.4.rev15 (Capella-Gutierrez, Silla-Martinez, & Gabaldon, 2009), and these processed loci were used for all subsequent downstream analyses. To help evaluate patterns in missing data or GC content in our taxa and UCE loci, the entire spruced dataset (consisting of 2,692 loci) was used as input for the program BaCoCa v1.1 (Kück & Struck, 2014).

### Matrix Generation

Seven datasets were generated using the PHYLUCE script phyluce_align_get_only_loci_with_min_taxa, differing by minimum taxon coverage (10%, 20%, 30%, 40%, 50%, 60%, & 70%). These datasets are summarized in Table 1. Each of these were concatenated into a combined matrix using the PHYLUCE script phyluce_align_format_nexus_files_for_raxml. The nucleotide substitution models were selected using ModelFinder (-m TEST; Kalyaanamoorthy *et al*., 2017) implemented in IQ-TREE v1.6.7 (Nguyen *et al*., 2015). The final 10% dataset was partitioned by SWSC-EN (v1.0.0; Tagliacollo & Lanfear, 2018) using default parameters, and ModelFinder was used to determine models for each partition.

**Table 1.** Statistics for the seven completeness datasets. The actual percentage of missing data (as determined by IQ-TREE), the number of characters in the concatenated alignment (as reported by IQ-TREE), the number of UCE loci, the number of parsimony informative sites in the concatenated alignment (as determined by IQ-TREE) and the percentage of parsimony informative sites (parsimony informative sites divided by total number of characters) are listed.

| completeness<br>matrix | % missing<br>data | characters | # of UCE<br>loci | parsimony<br>informative sites | parsimony<br>informative sites (%) |
| --- | --- | --- | --- | --- | --- |
| 10% | 68.54% | 893819 | 2508 | 249969 | 28.0 |
| 20% | 65.37% | 769977 | 2102 | 223184 | 29.0 |
| 30% | 60.12% | 576075 | 1481 | 174238 | 30.2 |
| 40% | 53.50% | 387311 | 968 | 118694 | 30.6 |
| 50% | 46.30% | 240305 | 593 | 76185 | 31.7 |
| 60% | 41.27% | 150330 | 349 | 48307 | 32.1 |
| 70% | 35.10% | 65338 | 146 | 21863 | 33.5 |

### Phylogenetic Analysis

Phylogenies of the seven concatenated datasets were inferred using Maximum Likelihood implemented in IQ-TREE (v1.6.7; Nguyen *et al*., 2015). Individual gene trees were also inferred using IQ-TREE for each dataset. Preliminary tests on the 50% dataset suggested that more thorough search parameters improved phylogenetic inference, especially for the gene trees. Bootstraps were replicated 1000 times (-bb 1000). One thousand initial parsimony trees were generated (-ninit 1000; default: 100) and the 100 top parsimony trees were used to initialize the candidate set (-ntop 100; default: 20), considering all possible nearest neighbor interchanges (-allnni; default: OFF). All 100 of these trees were maintained in the candidate set during the ML tree search (-nbest 100; default: 5) and unsuccessful runs were stopped after 1000 iterations (-nstop 1000; default: 100). The perturbation strength was set to 0.2 (-pers 0.2; default: 0.5), which is recommended for datasets with many short sequences. The gene trees from each of the seven datasets were provided as input for ASTRAL-III v5.6.2 (Zhang, Sayyari, & Mirarab, 2017) and analyzed using default settings, resulting in a species tree for each of the seven datasets. Trees were visualized in FigTree v1.4.3 (Rambaut, 2007).

### Alternative Measures of Support

Bootstrap values tend to overestimate nodal support on large datasets (e.g., Roycroft *et al*., 2019; Chan *et al*. 2020). We thus used Quartet Sampling v1.2 (Pease *et al*., 2018; https://www.github.com/fephyfofum/quartetsampling) and Concordance Factors analysis (Minh, Hahn, Lanfear 2020) implemented in IQ-TREE v1.7-beta12 to provide alternative support values. In order to investigate potential impacts of missing data on nodal support, the bootstrap, quartet concordance, gene concordance factor, and site concordance factor support values were recorded for each of the seven datasets differing in overall completeness.

Levels of support for each metric were divided into three categories: high, moderate, and low. The cutoffs for the nonparametric bootstrap (BS) follow those commonly used in the literature: high = 95-100%; moderate = 80-94%; low = < 80%. The cutoffs for Quartet Concordance (QC) were modified from Pease *et al*. (2018): high = 0.50-1; moderate = 0-.049; low = < 0. The formulation of cutoffs for gene concordance and site concordance factors (gCF and sCF, respectively) was less straightforward, as Minh, Hahn and Lanfear (2020) did not suggest what values constituted high or low support, except to say that sCF values below 33% may indicate problems at that node. To determine cutoffs for these concordance factors, support values were examined for uncontroversial, high-confidence nodes; those with QC≈1 and a preponderance of morphological data in support (e.g., congeneric taxa). The range of gCF and sCF values for these nodes were considered to be “high”. The same was then done for moderate support, with an alternative set of nodes generally well-supported by morphology but with only moderate QC support. This analysis suggested the following cutoffs for gCF: high = 70-100%; moderate = 20-69%; low = < 20%, and the following cutoffs for sCF: high = 60-100%; moderate = 33-59%; low = < 33%.

### Data Availability

Raw FASTQ files are deposited in the NCBI Sequence Read Archive (SRA), Bioproject accession number PRJNA666700. Specimen photographs, assembled contigs, processed alignments, BaCoCa output, scripts, tree files, and supplementary figures can be accessed via Figshare (DOI: 10.6084/m9.figshare.13140362).

## Results

### UCE Loci Recovery and Processing

Between 787 and 975 UCE loci were successfully extracted from the transcriptome assemblies (avg. 925), compared to the overall average of 944 loci. While there appeared to be no missing locus bias with transcriptome taxa, there was evidence of taxonomic bias in locus recovery. For example, an average of 1,169 and 1,130 loci were recovered for the subfamilies Asilinae and Laphriinae, respectively. Only an average of 629 loci, almost half as many, were recovered in the subfamily Dioctriinae. Using the 0.95 lognorm cutoff, Spruceup removed 538,970 sites from the concatenated alignment of all loci.

### Support & Missing Data

The topology for each % complete dataset became more similar as more loci (and concomitantly more missing data) were added (Figure 3). The topology stabilized (i.e., rf-distance ≈ zero) with 10-30% completeness for the concatenated analyses (Figure 3A) and 10-20% in the coalescent analyses (Figure 3B). However, the IQ-TREE and ASTRAL topologies were highly discordant (rf-distance > 40), and no amount of added loci was able to converge the topologies (Figure 3C).

**Fig 2.**
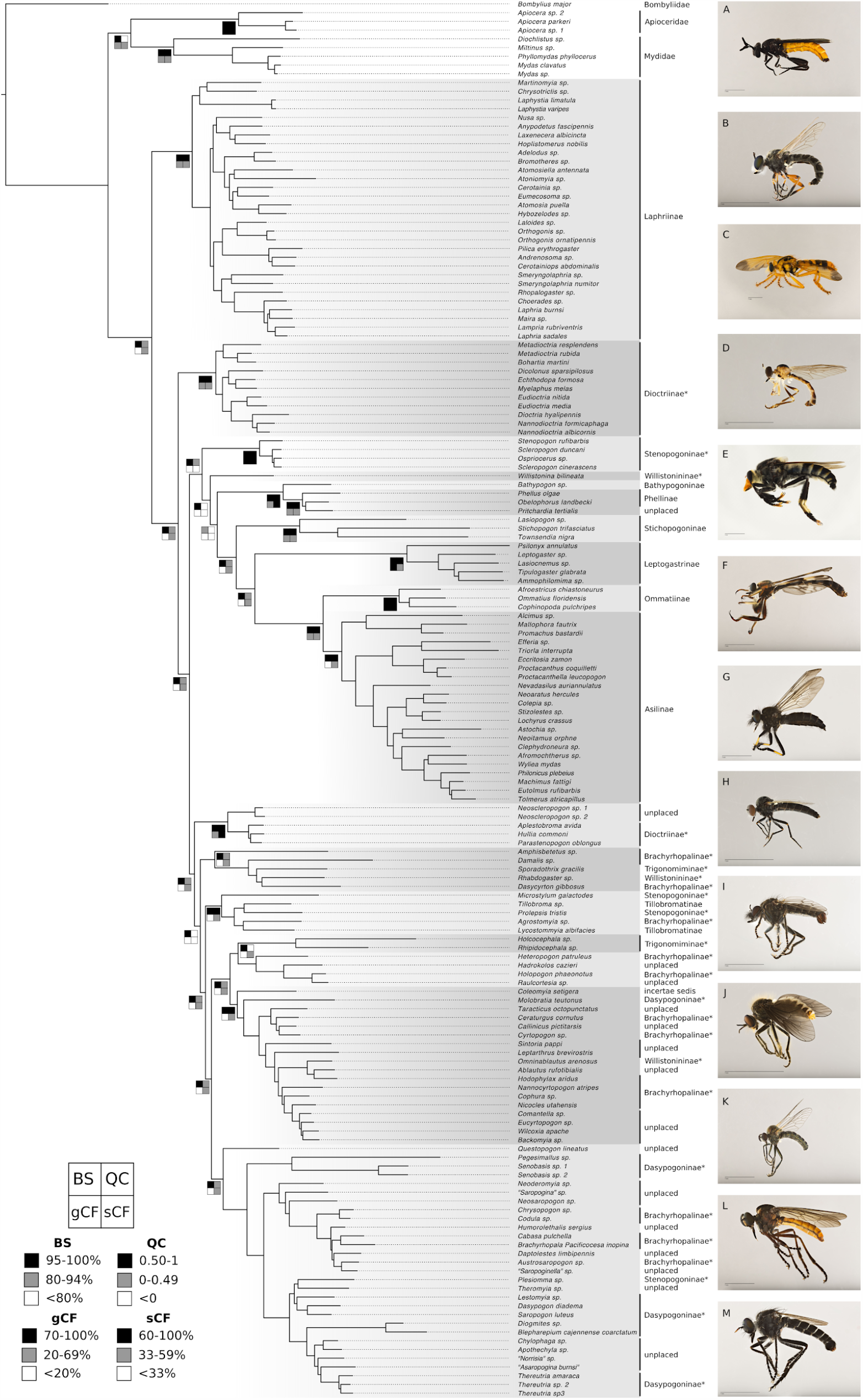
Maximum Likelihood tree of Asilidae inferred using IQ-TREE and the SWSC-EN partitioned 10% occupancy matrix. Major clades are depicted in alternating shades of grey to increase readability. The bootstrap (BS), quartet concordance (QC), gene concordance factor (gCF), and site concordance factor (sCF) support values are shown for subfamily and inter-subfamily nodes. Black equals high support, grey equals moderate support, and white equals low support (see legend for more details). The family or subfamily of each taxon is shown on the right, with those in a paraphyletic position denoted with an asterisk (*). Representative families and subfamilies are illustrated on the far right: (A) *Phyllomydas phyllocerus*, Mydidae; (B) *Laphystia sp*., Laphriinae; (C) *Laloides sp*., Laphriinae; (D) *Nannodioctria albicornis*, Dioctriinae; (E) *Phellus olgae*, Phellinae; (F) *Lasiocnemus sp*., Leptogastrinae; (G) *Stizolestes sp*., Asilinae; (H) *Amphisbetetus sp*., Brachyrhopalinae*; (I) *Tillobroma sp*., Tillobromatinae; (J) *Rhipidocephala sp*., Trigonomiminae; (K) *Sintoria pappi*, unplaced; (L) *Senobasis sp. 1*, Dasypogoninae; (M) *Thereutria sp. 2*, Dasypogoninae.

**Fig 3.**
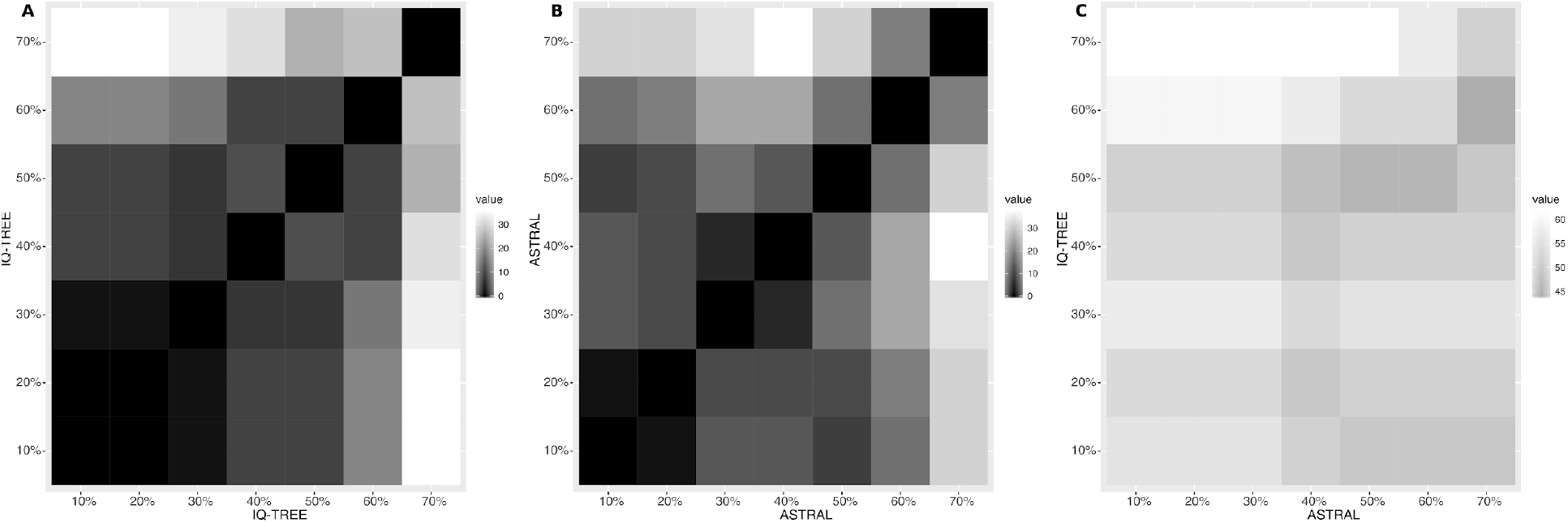
Robinson-Foulds distances of the seven % occupancy datasets for (A) IQ-TREE analyses, (B) ASTRAL analyses, and (C) IQ-TREE vs ASTRAL analyses.

Nodal support values across the entire topology, regardless of metric, did not vary greatly between % complete datasets (Figure 4). However, the different metrics each displayed characteristic behavior. Bootstrap values (BS) mostly reached maximum support, at or close to 100%. Quartet concordance values (QC) were predominantly high, but with a greater range of support (mostly 0.25-1). Gene concordance and site concordance factors likewise had a broad range of support, but these values were concentrated in the mid-to low-support end of the spectrum.

**Fig 4.**
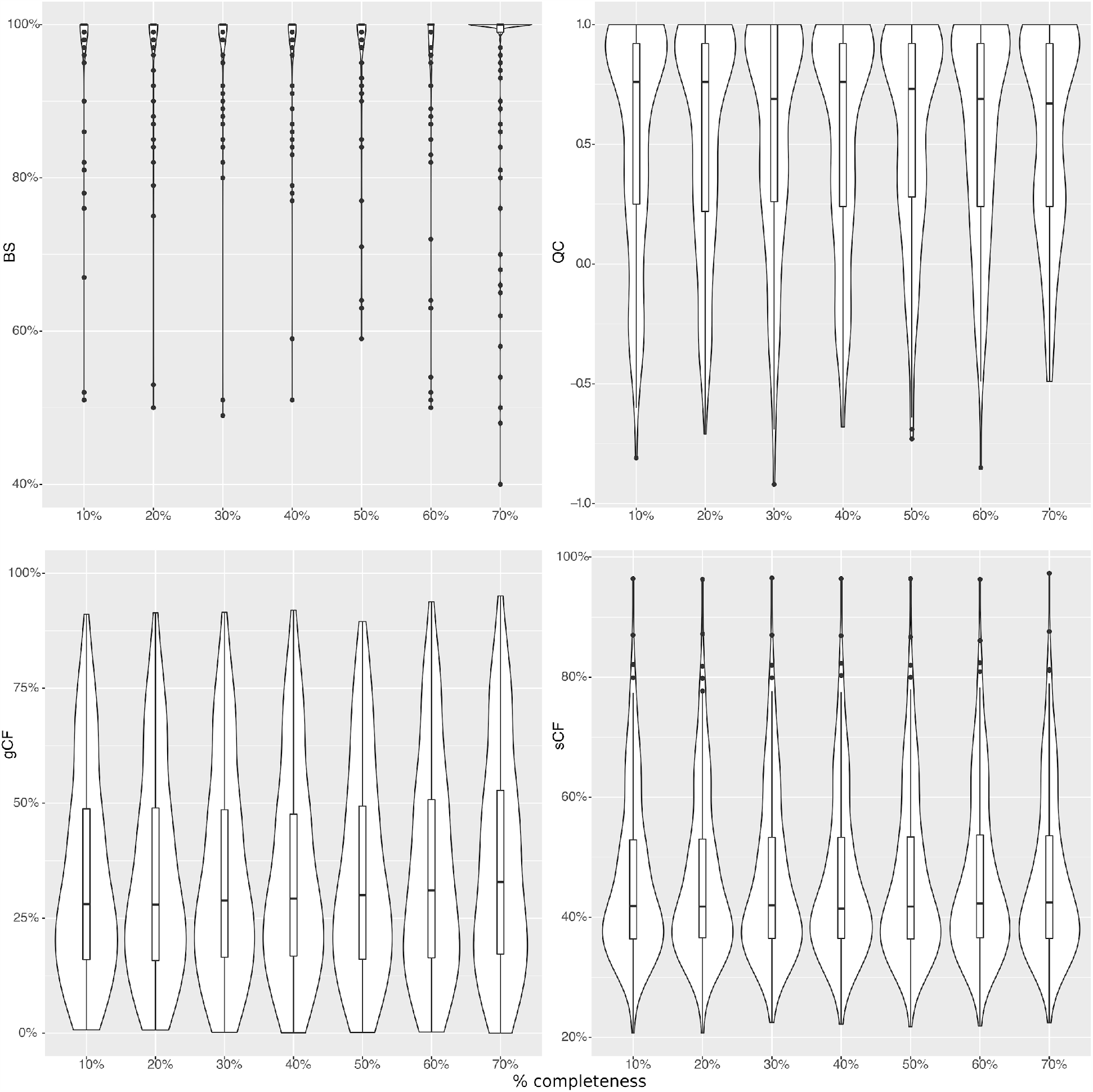
Violin plots and interquartile range of all nodal support values for each % occupancy matrix for the bootstrap (BS), quartet concordance (QC), gene concordance factor (gCF), and site concordance factor (sCF) metrics for the concatenated analyses.

### Phylogenetic Results

The prefered topology, based on the 10% matrix (the largest dataset) inferred with Maximum Likelihood in IQ-TREE, is shown in Figure 2. Only the subfamilies Laphriinae, Ommatiinae, Asilinae, Leptogastrinae, and Stichopogoninae were recovered as monophyletic *sensu* Dikow, 2009a. Phellinae was also recovered as monophyletic, with the addition of *Pritchardia* (currently unplaced *sensu* Dikow). Bathypogoninae had only one representative, so it was not possible to test the monophyly of this subfamily. The remaining subfamilies *sensu* Dikow (Brachyrhopalinae, Dasypogoninae, Dioctriinae, Stenopogoninae, Tillobromatinae, Trigonomiminae, Willistonininae) were recovered as paraphyletic or polyphyletic.

Apioceridae+Mydidae form a clade sister to a monophyletic Asilidae. The subfamily Laphriinae was recovered as sister to the rest of Asilidae, and Dioctriinae (in part) was found to be sister to a clade comprising two major lineages: the first contains Stenopogoninae (in part), Willistonininae (in part), Bathypogoninae, Phellinae, Stichopogoninae, Leptogastrinae, Ommatiinae, and Asilinae; the second clade contains Dioctriinae (in part), Trigonomiminae, Stenopogoninae (in part), Tillobromatinae, Brachyrhopalinae, and Dasypogoninae.

A representative ASTRAL tree inferred using the same 10% complete dataset is shown in Figure S1. Despite the high rf-distances between the ASTRAL and IQ-TREE topologies, at the subfamily level the trees are quite similar. Apioceridae+Mydidae is recovered as sister to Asilidae, Laphriinae is sister to the remaining Asilidae, and all the same subfamilies are paraphyletic. The other inter-subfamily relationships are largely the same between the two analyses, although the placement of Willistonininae and the clade containing *Damalis* occur in different positions. While the internal relationships differ in many places, the composition of the “subfamily-level” clades are otherwise identical to the concatenated analysis with two exceptions. *Holcocephala* is not recovered in a clade with *Rhipidocephala*, sister to Brachyrhopalinae (in part), but instead sister to the rest of Asilidae (after Laphriinae). Similarly, *Amphisbetetus* is not recovered as sister to the clade containing *Damalis*, but instead sister to the clade containing Stenopogoninae (in part), Tillobromatinae, Trigonomiminae (in part), Dasypogoninae, and Brachyrhopalinae.

## Discussion

### Transcriptomes in Arthropod UCE studies

It is now understood that many arthropod UCE loci occur in protein-coding regions, to a greater degree than UCEs found in vertebrates (Blaimer *et al*., 2019, Buenaventura *et al*. 2020; Hedin *et al*., 2019). The high number of UCE loci recovered here from asilid transcriptome assemblies, and the fact that they were recovered in phylogenetic positions with high support, lends further evidence to the idea that transcriptomes can be effectively combined with UCEs in phylogenomic analyses (Blaimer *et al*., 2019). However, it should be noted that UCE loci extracted from transcriptomes may be missing sequence data associated with genomic regions that are not translated, such as promoter regions or introns.

### Comparison of Measures of Support

The bootstrap is known to overestimate support in large datasets (e.g., Roycroft *et al*., 2019; Chan *et al*. 2020), which our UCE study corroborates. For example, most of the backbone nodes received maximum bootstrap support, but examination with alternative metrics revealed that support for many of these nodes was actually quite poor (Figure 2). Thus we conclude that bootstrap support should not be the sole metric reported for phylogenomic datasets.

Our results demonstrate that quartet concordance, gene concordance factors, and site concordance factors are more conservative and better suited to phylogenomic datasets (Figure 4). We find that gCF (and to a lesser extent sCF) tend to underestimate nodal support and are thus not an ideal replacement for the bootstrap. Instead, we recommend authors incorporate Quartet Concordance into their standard phylogenetic toolkit, as it strikes the best balance between the two extremes.

### Effect of Missing Data

Increasing matrix size (with a concomitant increase in missing data) had no significant effect on nodal support across the phylogeny, regardless of the metric used. These results have two implications. First, type I missing data (*sensu* Hosner *et al*., 2015; Sayyari *et al*., 2017), like that present in our dataset, may not have a detrimental effect on UCE phylogenomics. Indeed, numerous studies have shown that high amounts of missing data can have little to no effect on topology, at least for datasets with many taxa and/or loci (Wiens, 2003a; Wiens, 2003b; Driskell *et al*., 2004; Philippe *et al*., 2004; Fulton & Strobek, 2006; de la Torre-Barcena *et al*., 2009; Cho *et al*., 2011; Wiens & Morrill, 2011; Wiens & Tiu, 2012; Rubin *et al*., 2012; Roure *et al*., 2013; Jiang *et al*., 2014; Quek & Huang, 2019).

Second, nodal support is unaffected by matrix size or the amount of missing data. This result contradicts previous studies that found that bootstrap support increased with increasing matrix size, at least up to a certain threshold (Hosner *et al*., 2015; Streicher *et al*., 2016; Quek & Huang, 2019). We believe that this pattern can be explained by the bootstrap values becoming inflated as the matrix increases in size, and does not reflect genuine support. At the very least, there is no evidence from our particular dataset that support (from any metric) is correlated with matrix size. Minh, Hahn & Lanfear (2020) found the same result for their concordance factors. Authors have suggested that phylogenomic datasets with up to 40-70% missing data yield the best support and topology (Streicher *et al*., 2016; Quek & Huang, 2019). However, based on our results we believe that this is overly conservative - datasets with taxon occupancy as low as 10% (or possibly even lower) are just as well-supported as any other completeness matrix. We suggest that in the absence of other deciding factors, authors should prefer the topology inferred from the largest possible dataset while exploring changes across spectra of locus and taxon inclusion.

As noted in the results, there was evidence of taxonomic bias in missing data, with many fewer loci recovered in the subfamily Dioctriinae in particular. This bias may be due to genomic changes in this clade that resulted in the loss of loci (or modification of their ultraconserved cores) targeted by the Diptera-wide baitset. In contrast to those cited above, multiple studies have found that missing data, particularly when distributed non-randomly in loci or taxa, can mislead concatenation methods and some species tree methods (Agnarsson & May-Collado, 2008; Hartmann & Vision, 2008; Lemmon *et al*., 2009; Simmons, 2012a; Simmons, 2012b; Kvist & Siddall, 2013; Simmons, 2014; Xia, 2014; Xi *et al*., 2015; Sayyari *et al*., 2017). Furthermore, datasets with low taxon occupancy may result in topologically incongruent phylogenies with identical optimality scores (Sanderson *et al*., 2010; Steel and Sanderson, 2010; Sanderson *et al*., 2011). While it is not entirely clear what influences (if any) this bias in missing data had on our topology, the congruence of our phylogenetic results with a prior molecular study using an entirely different dataset (e.g., Dikow, 2009b) would seem to discount that idea.

### Discordance Between Concatenated and Coalescent Topologies

We find that the coalescent-based ASTRAL analyses performed suboptimally on our UCE datasets compared to IQ-TREE. The differing placements of *Holcocephala* and the clade containing *Damalis* are particularly noteworthy. *Holcocephala* and *Rhipidocephala* are members of the tribe Trigonomimini (Trigonomiminae), and their close relationship is generally well-supported by our concatenated ML analysis (BS=100; QC=0.86; gCF=35.3; sCF=45.5). The taxa also share numerous morphological synapomorphies such as: labella well developed and separated from prementum; anterior tentorial pits well developed, conspicuous, anteroventrally located; postpedicel cylindrical, same diameter throughout; prosternum laterally fused to proepisternum, with the former narrow above prothoracic coxa; dorsal (anterior) margin of prosternum with distinct flange-like projection; notopleural setae absent; postalar setae absent; apical scutellar setae absent; T9 and T10 entirely fused and indistinguishable; spurs on ovipositor absent (Dikow, 2009a).

The fact that both *Holcocephala* and the clade containing *Damalis* were recovered closer to the root of the tree by ASTRAL likely bears some significance, perhaps suggesting an underlying bias in the data. High GC content has been shown to sometimes negatively impact phylogenetic inference using UCEs (Bossert *et al*., 2017; Cruaud *et al*. 2020; but see Forthman, Miller & Kimball 2020). This is generally manifested as taxa clustering by similar GC content rather than by their “true” relationships. In our dataset, the bottom 35 taxa with the lowest % GC content are the outgroup taxa *Bombylius major* and most of the Mydidae (5/35; 0.328-0.367), nearly all Laphriinae (28/35; 0.345-0.374), and *Holcocephala* (0.354) and *Damalis* (0.363). *Rhipidocephala* by contrast has a % GC of 0.393 (avg. 0.402). In other words, the GC content of *Holcocephala* and *Damalis* is highly similar to the GC content of the outgroup taxa and the “basal” asilid subfamily Laphriinae. It is therefore plausible that GC bias played a role in the recovery of these two genera in positions close to the root of individual gene trees.

As a quick way to test this hypothesis, we removed all gene trees inferred from loci where *Holcocephala* had % GC below 0.35 (295/619 gene trees), creating a filtered dataset that was then provided as input to ASTRAL. This dataset induced numerous changes to the topology (rf-distance = 24), and, remarkably, recovered *Holcocephala* as sister to *Rhipidocephala* (Figure S2). It thus appears that low GC bias is in fact influencing phylogenetic inference of gene trees, and subsequently biasing the inferred topology of our summary coalescent method. We thus conclude that the ASTRAL topology does not reflect the “true” species tree, but rather is the result of shortcomings in our dataset: a misleading phylogenetic signal resulting from low GC bias, probably exacerbated by a combination of inadequate taxon sampling for these “trigonomiminine” lineages and by individual loci lacking sufficient phylogenetic signal. Removing the same 295 loci from the concatenated analysis results in a topology that is largely unchanged (rf-distance = 6). Our concatenated analysis thus appears to be more robust to both GC bias and loci removal.

### Phylogeny

Multiple deep relationships recovered in our topology agree with those found in previous studies. Apioceridae and Mydidae forming a clade sister to a monophyletic Asilidae has been recovered in many recent analyses (e.g., Dikow, 2009a, Dikow, 2009b, Dikow *et al*., 2017, Shin *et al*., 2018), although additional sampling of outgroup taxa from Asiloidea and lower Brachycera would be needed to confirm this. The position of Laphriinae as sister to the rest of Asilidae was recovered by the morphological analysis of Dikow (2009a). Finally, the relationship of (Stichopogoninae, (Leptogastrinae, (Ommatiinae, Asilinae))) was recovered by Dikow (2009b). The remaining inter-subfamily relationships in this study however appear to be novel.

Our phylogenetic analyses did not recover the subfamilies Brachyrhopalinae, Dasypogoninae, Dioctriinae, Stenopogoninae, Tillobromatinae, Trigonomiminae, or Willistonininae as monophyletic. The molecular-only topology of Dikow (2009b) also recovered these same subfamilies as paraphyletic, with the exception of Dioctriinae (the Australian members were not included). Instead, these Australian genera (*Aplestobroma, Hullia, Parastenopogon*) form a well-supported clade separate from Dioctriinae *sensu stricto*, and likely represent a new subfamily. One of the more remarkable findings is that the “goggle-eyed” flies (Trigonomiminae), characterized by wide, flattened heads, greatly enlarged eyes, and well-developed tentorial pits (such as *Damalis* and *Holcocephala*), do not form a natural group. Instead, we and Dikow (2009b) recover genera assigned to this subfamily in two separate and unrelated clades, meaning that this unusual condition has evolved at least twice in Asilidae.

The presence of an enlarged spur on the apex of the fore-tibia has long been considered an important character for recognizing the subfamily Dasypogoninae (e.g., Papavero, 1973). However, we recovered the unusual genus *Plesiomma* (Stenopogoninae: Plesiommatiini) in the Dasypogoninae (as did Dikow, 2009b), providing another example of the loss of the fore-tibial spur and the inadequacy of this character for subfamily delimitation. Consistent with recent phylogenetic analyses of Asilidae (Dikow, 2009a; Dikow, 2009b), we do not recover a monophyletic Dasypogoninae sensu Papavero, but instead recover genera with fore-tibial spurs in two separate clades: those allied to *Cyrtopogon* (Brachyrhopalinae), and those allied to *Dasypogon* (Dasypogoninae). However, we recover the tribes Brachyrhopalini (including *Brachyrhopala*, the type genus of Brachyrhopalinae) and Chrysopogonini (also Brachyrhopalinae) in the Dasypogoninae with high support. Because the latter is the older name, the subfamily Brachyrhopalinae will need to be discontinued, although this would leave the clade containing *Cyrtopogon* and allies without a subfamily name. These issues will need to be addressed in a future revision of the higher classification.

### Conclusion

Overall, this study demonstrates that the Diptera-wide UCE baitset can be successfully used to resolve family-level relationships in Diptera. Furthermore, it represents the largest and most comprehensive molecular phylogeny of Asilidae to date, allowing us to confidently state that half of the recognized subfamilies are not monophyletic and thus need to be reevaluated. A revised higher classification of Asilidae is in preparation that will incorporate this new understanding of subfamily relationships and composition (Cohen in prep). A refined classification will facilitate the systematic and evolutionary study of these diverse aerial predators.

## Supporting Information

**Table S1**. Specimen data. Collection locality, collection date, number of UCE loci recovered.

**Figure S1**. ASTRAL tree of the 10% completeness unfiltered dataset.

**Figure S2**. ASTRAL tree of the 10% completeness dataset, filtered to remove gene trees inferred from loci where *Holcocephala sp*. has % GC content of 0.35 or lower.

## Acknowledgements

We would like to thank Matt Bertone for helpful comments on the manuscript, and also the staff at the Australian National Insect Collection, the Queensland Museum, and Brigham Young University for the loan of specimens used in this study. A special thanks also goes to Darren Pollock for collecting many important specimens of Nearctic genera included in our phylogeny. CMC would like to thank Fred and Jean Hort for their generous hospitality while he conducted fieldwork in Western Australia, and Eric Fisher for assistance with several identifications. CMC also thanks Brant Faircloth and Michael Branstetter for help with troubleshooting library preparations. CMC was supported by the Graduate Research Fellowship Program (NSF-1645473) and fieldwork was funded by the AMNH Theodore Roosevelt Memorial Grant, the APS Lewis & Clark Fund for Exploration and Field Research, and the East Carolina University Department of Biology via startup funds provided to MSB.

## References

Abadi S, Azouri D, Pupko T, Mayrose I (2019) Model selection may not be a mandatory step for phylogeny reconstruction. Nature communications, 10, 934. doi:10.1038/s41467-019-08822-w.

Agnarsson I, May-Collado LJ (2008) The phylogeny of Cetartiodactyla: the importance of dense taxon sampling, missing data, and the remarkable promise of cytochrome b to provide reliable species-level phylogenies. Molecular phylogenetics and evolution, 48, 964–985. doi:10.1016/j.ympev.2008.05.046.

Baca SM, Alexander A, Gustafson GT, Short AEZ (2017) Ultraconserved elements show utility in phylogenetic inference of Adephaga (Coleoptera) and suggest paraphyly of ‘Hydradephaga’. Systematic entomology, 42, 786–795. doi:10.1111/syen.12244.

Barnes JK (2008) Review of the genus Ceraturgus Wiedemann (Diptera: Asilidae) in North America north of Mexico. Zootaxa, 1766, 1–45.

Blaimer BB, Brady SG, Schultz TR, Lloyd MW, Fisher BL, Ward PS (2015) Phylogenomic methods outperform traditional multi-locus approaches in resolving deep evolutionary history: a case study of formicine ants. BMC evolutionary biology, 15, 1–14. doi:10.1186/s12862-015-0552-5.

Blaimer BB, Lloyd MW, Guillory WX, Brady SG (2016) Sequence Capture and Phylogenetic Utility of Genomic Ultraconserved Elements Obtained from Pinned Insect Specimens. PloS one, 11, e0161531. doi:10.1371/journal.pone.0161531.

Borowiec ML (2019) Spruceup: fast and flexible identification, visualization, and removal of outliers from large multiple sequence alignments. Journal of Open Source Software, 4, 1635.

Bossert S, Murray EA, Almeida EAB, Brady SG, Blaimer BB, Danforth BN (2019) Combining transcriptomes and ultraconserved elements to illuminate the phylogeny of Apidae. Molecular phylogenetics and evolution, 130, 121–131. doi:10.1016/j.ympev.2018.10.012.

Branstetter MG, Longino JT, Ward PS, Faircloth BC (2017) Enriching the ant tree of life: enhanced UCE bait set for genome-scale phylogenetics of ants and other Hymenoptera (SPrice, Ed.). Methods in ecology and evolution/British Ecological Society, 8, 768–776. doi:10.1111/2041-210X.12742.

Buenaventura E, Lloyd MW,Perilla López JM, González VL, Thomas-Cabianca A, Dikow T (2020) Protein-encoding ultraconserved elements provide a new phylogenomic perspective of Oestroidea flies (Diptera: Calyptratae). Systematic entomology, 54, 1–23. doi:10.1111/syen.12443.

Bybee SM, Taylor SD, Riley Nelson C, Whiting MF (2004) A phylogeny of robber flies (Diptera: Asilidae) at the subfamilial level: molecular evidence. Molecular phylogenetics and evolution, 30, 789–797. doi:10.1016/S1055-7903(03)00253-7.

Capella-Gutiérrez S, Silla-Martínez JM, Gabaldón T (2009) trimAl: a tool for automated alignment trimming in large-scale phylogenetic analyses. Bioinformatics, 25, 1972–1973. doi:10.1093/bioinformatics/btp348.

Chan KO, Hutter CR, Wood PL, Grismer LL, Brown RM (2020) Larger, unfiltered datasets are more effective at resolving phylogenetic conflict: Introns, exons, and UCEs resolve ambiguities in Golden-backed frogs (Anura: Ranidae; genus Hylarana). Molecular phylogenetics and evolution, 151, 106899. doi:10.1016/j.ympev.2020.106899.

Cho S, Zwick A, Regier JC, Mitter C, Cummings MP, Yao J, Du Z, Zhao H, Kawahara AY, Weller S, Davis DR, Baixeras J, Brown JW, Parr C (2011) Can deliberately incomplete gene sample augmentation improve a phylogeny estimate for the advanced moths and butterflies (Hexapoda: Lepidoptera)? Systematic biology, 60, 782–796. doi:10.1093/sysbio/syr079.

Daniels G (1976) Three new species of Questopogon Dakin and Fordham (Diptera: Asilidae) from Australia. Proceedings of the Linnean Society of New South Wales Linnean Society of New South Wales, 100, 223–230.

Derrick J. Zwickl, David M. Hillis, Keith Crandall (2002) Increased Taxon Sampling Greatly Reduces Phylogenetic Error. Systematic biology, 51, 588–598. doi:10.1080/10635150290102339.

Dikow T (2009a) Phylogeny of Asilidae Inferred from Morphological Characters of Imagines (Insecta: Diptera: Brachycera: Asiloidea). Bulletin of the American Museum of Natural History, 2009, 1–175. doi:10.1206/603.1.

Dikow T (2009b) A phylogenetic hypothesis for Asilidae based on a total evidence analysis of morphological and DNA sequence data (Insecta: Diptera: Brachycera: Asiloidea). Organisms, diversity & evolution, 9, 165–188. doi:10.1016/j.ode.2009.02.004.

Dikow RB, Frandsen PB, Turcatel M, Dikow T (2017) Genomic and transcriptomic resources for assassin flies including the complete genome sequence of Proctacanthus coquilletti (Insecta: Diptera: Asilidae) and 16 representative transcriptomes. PeerJ, 5, e2951. doi:10.7717/peerj.2951.

Driskell AC, Ané C, Burleigh JG, McMahon MM, O’meara BC, Sanderson MJ (2004) Prospects for building the tree of life from large sequence databases. Science, 306, 1172–1174. doi:10.1126/science.1102036.

Faircloth BC (2013) Illumiprocessor: a trimmomatic wrapper for parallel adapter and quality trimming. Illumiprocessor: a trimmomatic wrapper for parallel adapter and quality trimming. doi:10.6079/J9ILL.

Faircloth BC (2017) Identifying conserved genomic elements and designing universal bait sets to enrich them (M Gilbert, Ed.). Methods in ecology and evolution / British Ecological Society, 8, 1103–1112. doi:10.1111/2041-210X.12754.

Faircloth BC, Branstetter MG, White ND, Brady SG (2015) Target enrichment of ultraconserved elements from arthropods provides a genomic perspective on relationships among Hymenoptera. Molecular ecology resources, 15, 489–501. doi:10.1111/1755-0998.12328.

Faircloth BC, McCormack JE, Crawford NG, Harvey MG, Brumfield RT, Glenn TC (2012) Ultraconserved Elements Anchor Thousands of Genetic Markers Spanning Multiple Evolutionary Timescales. Systematic biology, 61, 717–726. doi:10.1093/sysbio/sys004.

Fisher EM (2009) Asilidae (robber flies, assassin flies, moscas cazadoras, moscas ladronas). ‘Manual of Central American Diptera’. (Ed B.V. Brown, A. Borkent, J.M. Cumming, D.M. Wood, N.E. Woodley, M. Zumbado) pp. 585–632. (NRC Research Press).

Fulton TL, Strobeck C (2006) Molecular phylogeny of the Arctoidea (Carnivora): effect of missing data on supertree and supermatrix analyses of multiple gene data sets. Molecular phylogenetics and evolution, 41, 165–181. doi:10.1016/j.ympev.2006.05.025.

Gilbert PS, Chang J, Pan C, Sobel EM, Sinsheimer JS, Faircloth BC, Alfaro ME (2015) Genome-wide ultraconserved elements exhibit higher phylogenetic informativeness than traditional gene markers in percomorph fishes. Molecular phylogenetics and evolution, 92, 140–146. doi:10.1016/j.ympev.2015.05.027.

Glenn TC, Nilsen RA, Kieran TJ, Sanders JG, Bayona-Vásquez NJ, Finger JW, Pierson TW, Bentley KE, Hoffberg SL, Louha S, Garcia-De Leon FJ, Del Rio Portilla MA, Reed KD, Anderson JL, Meece JK, Aggrey SE, Rekaya R, Alabady M, Belanger M, Winker K, Faircloth BC (2019) Adapterama I: universal stubs and primers for 384 unique dual-indexed or 147,456 combinatorially-indexed Illumina libraries (iTru & iNext). PeerJ, 7, e7755. doi:10.7717/peerj.7755.

Grabherr MG, Haas BJ, Yassour M, Levin JZ, Thompson DA, Amit I, Adiconis X, Fan L, Raychowdhury R, Zeng Q, Chen Z, Mauceli E, Hacohen N, Gnirke A, Rhind N, di Palma F, Birren BW, Nusbaum C, Lindblad-Toh K, Friedman N, Regev A (2011) Full-length transcriptome assembly from RNA-Seq data without a reference genome. Nature biotechnology, 29, 644–652. doi:10.1038/nbt.1883.

Hartmann S, Vision TJ (2008) Using ESTs for phylogenomics: can one accurately infer a phylogenetic tree from a gappy alignment? BMC evolutionary biology, 8, 95. doi:10.1186/1471-2148-8-95.

Harvey MG, Smith BT, Glenn TC, Faircloth BC, Brumfield RT (2016) Sequence Capture versus Restriction Site Associated DNA Sequencing for Shallow Systematics. Systematic biology, 65, 910–924. doi:10.1093/sysbio/syw036.

Heath TA, Hedtke SM, Hillis DM (2008) Taxon sampling and the accuracy of phylogenetic analyses. Journal of systematics and evolution, 46, 239–257.

Hedin M, Derkarabetian S, Alfaro A, Ramírez MJ, Bond JE (2019) Phylogenomic analysis and revised classification of atypoid mygalomorph spiders (Araneae, Mygalomorphae), with notes on arachnid ultraconserved element loci. PeerJ, 7, e6864. doi:10.7717/peerj.6864.

Hillis DM (1998) Taxonomic sampling, phylogenetic accuracy, and investigator bias. Systematic biology, 47, 3–8. doi:10.1080/106351598260987.

Hosner PA, Faircloth BC, Glenn TC, Braun EL, Kimball RT (2016) Avoiding Missing Data Biases in Phylogenomic Inference: An Empirical Study in the Landfowl (Aves: Galliformes). Molecular biology and evolution, 33, 1110–1125. doi:10.1093/molbev/msv347.

Jiang W, Chen S-Y, Wang H, Li D-Z, Wiens JJ (2014) Should genes with missing data be excluded from phylogenetic analyses? Molecular phylogenetics and evolution, 80, 308–318. doi:10.1016/j.ympev.2014.08.006.

Kalyaanamoorthy S, Minh BQ, Wong TKF, von Haeseler A, Jermiin LS (2017) ModelFinder: fast model selection for accurate phylogenetic estimates. Nature Methods, 14, 587–589. doi:10.1038/nmeth.4285.

Karl E (1959) Vergleichend-morphologische Untersuchungen der männlichen Kopulationsorgane bei Asiliden. Beitrage zur Entomologie/Deutches Entomologisches Institut, 9, 619–680.

Kück P, Struck TH (2014) BaCoCa--A heuristic software tool for the parallel assessment of sequence biases in hundreds of gene and taxon partitions. Molecular phylogenetics and evolution, 70, 94–98.

Kvist S, Siddall ME (2013) Phylogenomics of Annelida revisited: a cladistic approach using genome-wide expressed sequence tag data mining and examining the effects of missing data. Cladistics, 29, 435–448.

Lemmon AR, Brown JM, Stanger-Hall K, Lemmon EM (2009) The effect of ambiguous data on phylogenetic estimates obtained by maximum likelihood and Bayesian inference. Systematic biology, 58, 130–145. doi:10.1093/sysbio/syp017.

Londt JGH, Dikow T (2017) Asilidae. ‘Manual of Afrotropical Diptera. Volume 2. Nematocerous Diptera and lower Brachycera’. (Ed A.H. Kirk-Spriggs & B.J. Sinclair) pp. 1097–1182. (South African National Biodiversity Institute).

Manthey JD, Campillo LC, Burns KJ, Moyle RG (2016) Comparison of Target-Capture and Restriction-Site Associated DNA Sequencing for Phylogenomics: A Test in Cardinalid Tanagers (Aves, Genus: Piranga). Systematic biology, 65, 640–650. doi:10.1093/sysbio/syw005.

Martin CH (1959) The Holopogon Complex of North America, Excluding Mexico, with the Descriptions of a New Genus and a New Subgenus (Diptera, Asilidae). American Museum novitates, 1980, 1–40.

Minh BQ, Hahn MW, Lanfear R (2020) New Methods to Calculate Concordance Factors for Phylogenomic Datasets. Molecular biology and evolution, 37, 2727–2733. doi:10.1093/molbev/msaa106.

Nguyen L-T, Schmidt HA, von Haeseler A, Minh BQ (2015) IQ-TREE: a fast and effective stochastic algorithm for estimating maximum-likelihood phylogenies. Molecular biology and evolution, 32, 268–274. doi:10.1093/molbev/msu300.

Papavero N (1973) Studies of Asilidae (Diptera) Systematics and Evolution. I. A Preliminary Classification in Subfamilies. Arquivos de Zoologia, 23, 217–274.

Papavero N, Artigas JN, Lamas CJE (2013) Manual of Neotropical Diptera. Asilidae. Neotropical Diptera, 18, 1–320.

Paramonov SJ (1958) A Review of Australian Species of Laphria (Asilidae, Diptera), with Descriptions of Three New Species from Lord Howe Island. Pacific science, 12, 92–105.

Pease JB, Brown JW, Walker JF, Hinchliff CE, Smith SA (2018) Quartet Sampling distinguishes lack of support from conflicting support in the green plant tree of life. American journal of botany, 105, 385–403. doi:10.1002/ajb2.1016.

Philippe H, Snell EA, Bapteste E, Lopez P, Holland PWH, Casane D (2004) Phylogenomics of eukaryotes: impact of missing data on large alignments. Molecular biology and evolution, 21, 1740–1752. doi:10.1093/molbev/msh182.

Quek ZBR, Huang D (2019) Effects of missing data and data type on phylotranscriptomic analysis of stony corals (Cnidaria: Anthozoa: Scleractinia). Molecular phylogenetics and evolution, 134, 12–23. doi:10.1016/j.ympev.2019.01.012.

Rambaut A (2016) FigTree v1.4.3. http://tree.bio.ed.ac.uk/software/figtree/.

Rhodén C, Wahlberg E (2020) The phylogeny of Empis and Rhamphomyia (Diptera, Empididae) investigated using UCEs including an over 150 years old museum specimen. Evolutionary Systematics, 4, 21.

Roure B, Baurain D, Philippe H (2013) Impact of missing data on phylogenies inferred from empirical phylogenomic data sets. Molecular biology and evolution, 30, 197–214. doi:10.1093/molbev/mss208.

Roycroft EJ, Moussalli A, Rowe KC (2020) Phylogenomics Uncovers Confidence and Conflict in the Rapid Radiation of Australo-Papuan Rodents. Systematic biology, 69, 431–444. doi:10.1093/sysbio/syz044.

Rubin BER, Ree RH, Moreau CS (2012) Inferring phylogenies from RAD sequence data. PloS one, 7, e33394. doi:10.1371/journal.pone.0033394.

Sanderson MJ, McMahon MM, Steel M (2010) Phylogenomics with incomplete taxon coverage: the limits to inference. BMC evolutionary biology, 10, 155. doi:10.1186/1471-2148-10-155.

Sanderson MJ, McMahon MM, Steel M (2011) Terraces in phylogenetic tree space. Science, 333, 448–450. doi:10.1126/science.1206357.

Sayyari E, Whitfield JB, Mirarab S (2017) Fragmentary Gene Sequences Negatively Impact Gene Tree and Species Tree Reconstruction. Molecular biology and evolution, 34, 3279–3291. doi:10.1093/molbev/msx261.

Shin S, Bayless KM, Winterton SL, Dikow T, Lessard BD, Yeates DK, Wiegmann BM, Trautwein MD (2018) Taxon sampling to address an ancient rapid radiation: a supermatrix phylogeny of early brachyceran flies (Diptera). Systematic entomology, 43, 277–289. doi:10.1111/syen.12275.

Simmons MP (2012a) Misleading results of likelihood-based phylogenetic analyses in the presence of missing data. Cladistics, 28, 208–222. https://onlinelibrary.wiley.com/doi/abs/10.1111/j.1096-0031.2011.00375.x.

Simmons MP (2012b) Radical instability and spurious branch support by likelihood when applied to matrices with non-random distributions of missing data. Molecular phylogenetics and evolution, 62, 472–484. doi:10.1016/j.ympev.2011.10.017.

Simmons MP (2014) A confounding effect of missing data on character conflict in maximum likelihood and Bayesian MCMC phylogenetic analyses. Molecular phylogenetics and evolution, 80, 267–280. doi:10.1016/j.ympev.2014.08.021.

Smith BT, Harvey MG, Faircloth BC, Glenn TC, Brumfield RT (2014) Target Capture and Massively Parallel Sequencing of Ultraconserved Elements for Comparative Studies at Shallow Evolutionary Time Scales. Systematic biology, 63, 83–95. doi:10.1093/sysbio/syt061.

Springer MS, Gatesy J (2018) On the importance of homology in the age of phylogenomics. Systematics and biodiversity 16, 210–228. doi:10.1080/14772000.2017.1401016.

Starrett J, Derkarabetian S, Hedin M, Bryson RW Jr, McCormack JE, Faircloth BC (2017) High phylogenetic utility of an ultraconserved element probe set designed for Arachnida. Molecular ecology resources, 17, 812–823. doi:10.1111/1755-0998.12621.

Steel M, Sanderson MJ (2010) Characterizing phylogenetically decisive taxon coverage. Applied mathematics letters, 23, 82–86. doi:10.1016/j.aml.2009.08.009.

Streicher JW, Schulte JA 2nd, Wiens JJ (2016) How Should Genes and Taxa be Sampled for Phylogenomic Analyses with Missing Data? An Empirical Study in Iguanian Lizards. Systematic biology, 65, 128–145. doi:10.1093/sysbio/syv058.

Tagliacollo VA, Lanfear R (2018) Estimating Improved Partitioning Schemes for Ultraconserved Elements. Molecular biology and evolution, 35, 1798–1811. doi:10.1093/molbev/msy069.

Torre-Bárcena JE de la, de la Torre-Bárcena JE, Kolokotronis S-O, Lee EK, Stevenson DW, Brenner ED, Katari MS, Coruzzi GM, DeSalle R (2009) The Impact of Outgroup Choice and Missing Data on Major Seed Plant Phylogenetics Using Genome-Wide EST Data. PLoS one, 4, e5764. doi:10.1371/journal.pone.0005764.

Van Dam MH, Lam AW, Sagata K, Gewa B, Laufa R, Balke M, Faircloth BC, Riedel A (2017) Ultraconserved elements (UCEs) resolve the phylogeny of Australasian smurf-weevils. PloS one, 12, e0188044. doi:10.1371/journal.pone.0188044.

Wiegmann BM, Trautwein MD, Winkler IS, Barr NB, Kim J-W, Lambkin C, Bertone MA, Cassel BK, Bayless KM, Heimberg AM, Wheeler BM, Peterson KJ, Pape T, Sinclair BJ, Skevington JH, Blagoderov V, Caravas J, Kutty SN, Schmidt-Ott U, Kampmeier GE, Thompson FC, Grimaldi DA, Beckenbach AT, Courtney GW, Friedrich M, Meier R, Yeates DK (2011) Episodic radiations in the fly tree of life. Proceedings of the National Academy of Sciences of the United States of America 108, 5690–5695. doi:10.1073/pnas.1012675108.

Wiens JJ (2003a) Incomplete taxa, incomplete characters, and phylogenetic accuracy: is there a missing data problem? Journal of Vertebrate Paleontology, 23, 297–310. doi:10.1671/0272-4634(2003)023[0297:ITICAP]2.0.CO;2.

Wiens JJ (2003b) Missing data, incomplete taxa, and phylogenetic accuracy. Systematic biology, 52, 528–538. doi:10.1080/10635150390218330.

Wiens JJ, Morrill MC (2011) Missing data in phylogenetic analysis: reconciling results from simulations and empirical data. Systematic biology, 60, 719–731. doi:10.1093/sysbio/syr025.

Wiens JJ, Tiu J (2012) Highly incomplete taxa can rescue phylogenetic analyses from the negative impacts of limited taxon sampling. PloS one, 7, e42925. doi:10.1371/journal.pone.0042925.

Wood GC (1981) Asilidae. ‘Manual of Nearctic Diptera Volume 1’. (Ed J.F. McAlpine, B.V. Peterson, G.E. Shewell, H.J. Teskey, J.R. Vockeroth, D.M. Wood) pp. 549–573. (Research Branch Agriculture Canada).

Xi Z, Liu L, Davis CC (2016) The Impact of Missing Data on Species Tree Estimation. Molecular biology and evolution, 33, 838–860. doi:10.1093/molbev/msv266.

Xia X (2014) Phylogenetic Bias in the Likelihood Method Caused by Missing Data Coupled with Among-Site Rate Variation: An Analytical Approach. In ‘Bioinformatics Research and Applications’, 12–23. (Springer International Publishing) doi:10.1007/978-3-319-08171-7_2.

Zhang C, Sayyari E, Mirarab S (2017) ASTRAL-III: Increased Scalability and Impacts of Contracting Low Support Branches. ‘Comparative Genomics’. (Eds J Meidanis, L Nakhleh) Lecture Notes in Computer Science. pp. 53–75. (Springer International Publishing: Cham) doi:10.1007/978-3-319-67979-2_4.

Zhang YM, Williams JL, Lucky A (2019) Understanding UCEs: A Comprehensive Primer on Using Ultraconserved Elements for Arthropod Phylogenomics. Insect Systematics and Diversity, 3. doi:10.1093/isd/ixz016.

